# Barcoding analysis of HIV drug resistance mutations using Oxford Nanopore MinION (ONT) sequencing

**DOI:** 10.1101/240077

**Authors:** Claudia Gonzalez, Jessica Gondola, Alma Y Ortiz, Juan M Castillo, Juan M Pascale, Alexander A Martinez

## Abstract

Determination of HIV drug resistance (HIVDR) is becoming an integral baseline HIV evaluation for newly infected subjects, as the level of pre-treatment resistance is increasing worldwide. Until now, the gold standard for monitoring ART mutations is the Sanger sequencing method, however, next-generation sequencing technologies (NGS) because high-throughput capability, are gaining attention as a method for detection of HIVDR. In the present work, we evaluated the use of the Oxford Nanopore Technologies (ONT) MinION as an alternative method for detection of drug resistance mutations in pre-treatment HIV positive subjects.

We evaluate 36 samples taken during November 2016 from treatment naïve subjects with age greater than 18 years old, who went to the lab for their first HIV monitoring. To evaluate the agreement between Sanger and MinION generated sequences, we aligned the sequences (∼1200bp) with muscle v. 3.8.31. Then we counted the differences and calculated the p-distance of the obtained sequences, comparing paired sequences and grouping Sanger and MinION obtained sequences. The percentage of similarity among each sequence was also evaluated.

All samples were submitted to the Standford University HIV drug resistance database (HIVdb version 8.4). Then we compared the resistance predictions obtained from the sequences generated by Sanger and MinION methods.

Results: The median of available pores was 1314 for the first run, 1215 for the second run, and 536 for the third run. After 3 hours with SQK-NSK007 a total of 18803 2D reads were base-called and in 16577 reads (88%) a barcode was detected.

Comparing the nucleotide differences of each sample, we observed that 23 (74%) samples had identical sequence, for the other samples the percentage of identity among each analyzed sequence was greater than 95%. A good positive predictive value (100%) in the estimation of drug resistance mutations in the groups of protease inhibitors (PI), nucleoside reverse transcriptase inhibitors (NRTIs), and non-nucleoside reverse transcriptase inhibitors (NNRTIs).

We present an approach for the analysis of HIV reads generated with MinION ONT, further studies are guaranteed before the application of this methodology in clinical settings to assess its suitability for HIVDR testing.

## Introduction

The HIV replication cycle is characterized by the incorporation of nucleotide substitutions because an error prone polymerase (1), that ultimately in presence of antiretroviral therapy (ART) pressure, allows the selection of mutant strains, that confer resistance to treatment (2). The gold standard for the determination of ART mutations is the Sanger sequencing methodology, however, next-generation sequencing (NGS) as the Illumina technology is gaining attention for their high-throughput capability, as a method for determination of ART in HIV (3). With the Illumina-based NGS method, samples can be processed in pools, reducing costs of pre-treatment HIV drug resistance surveillance programs. Nevertheless, a third-generation sequencing technology (TGS) ONT - MinION is arising in popularity within the scientific community (4, 5). This method can also be performed in pools and is based in the sequencing of single DNA molecules. By using a solid-state graphene membrane, coated with proteins forming nano-pore channels, this method allows the DNA to pass through the pore, creating a characteristic disruption in current which is sensed by an array of sensor chips (6). However, it has been reported a low level of accuracy of the reads generated with ONT, this could be a serious caveat in case of sequencing a high mutation rate virus as HIV. In the present article, we evaluated the capability of ONT for the determination of pre-treatment HIV drug resistance PHIVDR and results were compared with the gold-standard method.

## Methodology

### Ethic Statement

Samples from an ongoing pre-treatment HIV drug surveillance study were included. This study was approved by the Gorgas Memorial Institutional Bioethics Review Board (GMIBRB). All the participant signed an informed consent for the use of their sample in the evaluation of HIV resistance. For this study, samples were decoded and analyzed anonymously.

### Sample collection

In this study, 36 samples were taken during November 2016 from treatment naïve subjects with age greater than 18 years old, who went to the genomics and proteomics laboratory of the Gorgas Memorial Institute for their first monitoring.

### RNA extraction, amplification

HIV-1 pol region sequences were obtained using an “in-house” drug resistance genotyping method (described below). Plasma samples were centrifuged at 20,000 *G* 1 hour and viral RNA was extracted using the QIAamp Viral RNA Mini kit (Qiagen Inc., Valencia, CA). Reverse transcription was performed using the Thermoscript Reverse Transcriptase enzyme (Invitrogen, Carlsbad, CA), following the manufacturer’s instructions. A 1.2 kb fragment of the HIV-1 pol gene spanning the complete protease (Pro, codons 1-99) and part of reverse transcriptase (RT, codons 1-235) was amplified by a nested polymerase chain reaction (PCR) using Platinum Taq polymerase (Invitrogen, Carlsbad, CA). Both PCR reactions were performed in a final volume of 50 μL with 1.8mM MgCl2, 0.2mM dNTP mix, 0.2 μM each primer (7). The first-round PCR was carried out under the following conditions: 94°C, 2min, 30 cycles at 94°C-20s, 50°C-20s, 72°C-90s, final extension of 72°C-6min. Second round PCR conditions were: 94°C, 2min, 40 cycles at 94°C −20s, 50^°^C-20s, 72°C-90s, final extension of 72°C-6min. PCR products were electrophoresed on 1% agarose gels and DNA bands of expected size purified using PCR purification kit (Qiagen Inc., Valencia, CA).

### Sanger sequencing

Direct cycle sequencing was performed with seven overlapping segment primers using the ABI Prism BigDye Terminator v3.1 Cycle Sequencing kit and an ABI PRISM 3130xl Genetic Analyzer (Life Technologies, Carlsbad, CA). Primers used for PCR and sequencing have been previously described (7, 8). Sequence fragments were assembled using the Sequencher software, version 4.5 (GeneCodes, Ann Arbor, MI).

### MinION library preparation

This work was completed as part of the Oxford Nanopore Technologies (ONT) MinION early-access program. The PCR products were prepared according to the ONT sequencing kits (SQK-NSK007) for one run and the ligation sequencing kit 1D (SQK-LSK108) for two additional runs.

Briefly, 1 μg of each PCR product in 45 μL of molecular grade water (Promega), was used for the end repair process with NEBNext Ultra II End-repair/dA-tailing (New England Biolabs, Ipswich, MA, USA), adding 7μL of End-prep buffer and 3 μL of enzyme mix. The mix was incubated at 20^°^C and 5 minutes at 65 ^°^C using a thermal cycler (Applied). Then the end repaired DNA was purified using 1:1 Agencourt AMPure XP beads (Beckman Coulter Inc. Pasadena CA, USA), and eluted in 31μL (25μL for SQK-μLSK108) of molecular grade water. The recovery was evaluated with the Qubit dsDNA BR Assay Kit. As the fragment end-repaired had less than 3 kb, the amount of DNA was adjusted to use 0.2 pmoles per end repaired fragment. For the barcode ligation 22.5 μL (∼500ng) of the end-prepped fragments were ligated with 2.5 μL of one of the barcode (NB01-NB12) of the native barcoding kit (EXP-NBD002 for SQK-NSK007 and EXP-NBD103 for SQK-μLSK108) using 25 μL of Blunt/TA ligase MM (New England Biolabs, Ipswich, MA, USA). After 10 min of incubation at room temperature, 50μL of Agencourt AMPure XP beads was added for a purification process, as per manufactured indications. The barcoded ligated amplicon was eluted in 26μL of molecular grade water. After an equimolar amount of pooled barcoded amplicons (∼700 ng in 38μL) was mixed, for the SQK-NSK007 with 10μL of native barcoding adapter mix (BAM), 2μL of native barcoding hairpin adaptor (BHP) and 150μL Blunt/TA ligase. Next the mix was incubated for 10 min, at room temperature (∼25°C), following 1μL of Tether was added and incubated for additional 10 min. The adapted and tethered library was purified using MyOne C1 beads (Invitrogen), as per manufactured recommendations. For the SQK-LSK108 an equimolar amount of pooled barcoded amplicons (∼700 ng in 38μL) was mixed with 12μL of native barcoding adapter mix (BAM), and 50μL Blunt/TA ligase. Next the mix was incubated for 10 min, at room temperature (∼25°C), following 40μL of AMpure XP beads was added and incubated at RT for 5 min in a rotator. Then 140μL of adapter bead binding buffer (ABB) was added and the adapted library pellet on magnet, finally, the adapted library was eluted in 15μL of elution buffer (ELB), this is the pre-sequencing mix.

### MinION

The sequencing library was loaded into the flow cell, as per manufactured indication and ran for 3 hours with the 48-H sequencing protocol on the MinKNOW software (version: 1.2.8.0).

### MinION Data Analysis

Raw sequence data was uploaded and base-called with the cloud-based Metrichor workflow 2D Basecalling plus Barcoding for FLO-MIN105 250bps for SQK-NSK007 kits. For LSK-NSK108 kits, reads were base-called with albacore (v. 1.2.2). Reads were extracted from fast5 format in fastqc and fasta with nanopolish extract (v.0.6.1). A consensus draft of the reads was generated with canu 1.3, using the following contigFilter options in command line: 1 1200 1.0 1.0 2, then the draft consensus was mapped with LASTAL (v. 759) with the command line: -s 2 –q 1 –b 1 –a 1 –e 45 –T 0 –Q 0 –a 1. Samtools was used to convert the data in bam format. Following the assembly was improved with nanopolish eventaling (v.0.6.1) and the consensus generated with nanopolish variants. Pipeline used in this study is archive at https://github.com/AAMCgenomics/hivminion. The sanger sequences obtained for the 31 samples analyzed in this study were deposited at the GenBank database, under accession numbers MG982909 to MG982931. The ONT reads were uploaded to European nucleotide archive (ENA EMBL-EBI) under accession numbers ERR2318840 to ERR2318870.

### Comparative Analysis among Sanger and MinION generated sequences

To evaluate the agreement between Sanger and MinION generated sequences, we aligned the sequences (∼1200bp) with muscle v. 3.8.31. Then we counted the differences and calculated the p-distance of the obtained sequences, comparing paired sequences and grouping Sanger and MinION obtained sequences. The percentage of similarity among each sequence was also evaluated.

All samples were submitted to the Standford University HIV drug resistance database (HIVdb version 8.4), to evaluate the impact of the mutations observed in the obtained sequences. Then we compared the resistance predictions obtained from the sequences generated by Sanger and MinION methods.

## Results

Three runs were performed, one of them with the SQK-NSK007 kit and two with the LSK-NSK108 kit. From the 36 samples collected the complete sequence was not obtained in Sanger for 4 samples and one sample was evaporated before adding to the MinION sequencer, therefore, 31 samples were included in three runs. The median of available pores was 1314 for the first run, 1215 for the second run, and 536 for the third run. After 3 hours with SQK-NSK007 a total of 18803 2D reads were base-called and in 16577 reads (88%) a barcode was detected. The second run performed with the kit LSK-NSK108 which yields a total of 42138 base-called reads and in 38764 (91.9 %) a barcode was detected. For the third run after 6 hours with LSK-NSK108 64220 reads were base-called, in 41335 (64.4%) of them a barcode was detected. Supplementary Table 1 shows the number of total reads and nucleotide length statistics of each barcode for the three runs.

**Table 1.** Nucleotide differences and distance between the Sanger and MiION obtained sequences (n=31) and the samples itself. The nucleotide distance within both method and 31 samples analyzed was calculated with p-distance as implemented in MEGA 7.0 ^*^The completed sequence was not obtained in Sanger for 4 sequences and one sample was evaporated before start the MinION reaction.

| group | N° of nucleotide differences (SE) | % of similarity | nucleotide distance (SE) |
| --- | --- | --- | --- |
| Sanger | ----- | ----- | ----- |
| MinION | 49.6 (3.3) | 95.14 | 0.1 (0) |
| 10278 | 0 (0) | 100 | 0 (0) |
| 10986 | 1 (0.96) | 99.9 | 0 (0) |
| 11133 | 0 (0) | 100 | 0 (0) |
| 11483 | 3 (1.62) | 99.7 | 0 (0) |
| 12967 | 25 (4.99) | 97.5 | 0.03 (0.01) |
| 14023 | 0 (0) | 100 | 0 (0) |
| 15213 | 0 (0) | 100 | 0 (0) |
| 15214 | 0 (0) | 100 | 0 (0) |
| 15301 | 0 (0) | 100 | 0 (0) |
| 15492 | 0 (0) | 100 | 0 (0) |
| 15606 | 0 (0) | 100 | 0 (0) |
| 16714 | 1 (0.94) | 99.9 | 0 (0) |
| 16715 | 0 (0) | 100 | 0 (0) |
| 16928 | 0 (0) | 100 | 0 (0) |
| 17157 | 0 (0) | 100 | 0 (0) |
| 17237 | 0 (0) | 100 | 0 (0) |
| 17461 | 0 (0) | 100 | 0 (0) |
| 18127 | 0 (0) | 100 | 0 (0) |
| 18130 | 12 (3.32) | 99.9 | 0.01 (0) |
| 2337 | 53 (7.5) | 94.7 | 0.06 (0.01) |
| 3463 | 0 (0) | 100 | 0 (0) |
| 3467 | 0 (0) | 100 | 0 (0) |
| 3841 | 3 (1.67) | 99.7 | 0 (0) |
| 3988 | 0 (0) | 100 | 0 (0) |
| 4125 | 0 (0) | 100 | 0 (0) |
| 4175 | 0 (0) | 100 | 0 (0) |
| 5442 | 1 (0.97) | 99.9 | 0 (0) |
| 6154 | 0 (0) | 100 | 0 (0) |
| 8600 | 0 (0) | 100 | 0 (0) |
| 9254 | 0 (0) | 100 | 0 (0) |
| 9683 | 0 (0) | 100 | 0 (0) |

The median of the length of the reads obtained was according with the input amplicon length, this indicates that there was not fragmentation during the library preparation (Supplementary Table 1).

Comparing the nucleotide differences of each sample, we observed that 23 (74%) samples had identical sequence, for the other samples the percentage of identity among each analyzed sequence was greater than 95%. When comparing the nucleotide distance, all sequences had 0.1 or less nucleotide distance among them, table #1.

Mutations associated to drug resistance were found in 12 samples. The following mutations were detected: M41L, K103N, E138A, V179E, V179D, V82L, Q58E (Supplementary table 2). Among the polymorphisms observed, there were some differences in the number detected by MinION vs Sanger. Most are attributable to the fact that with MinION we can sequence the whole fragment, covering the 5’-3’ends.

**Table 2.**
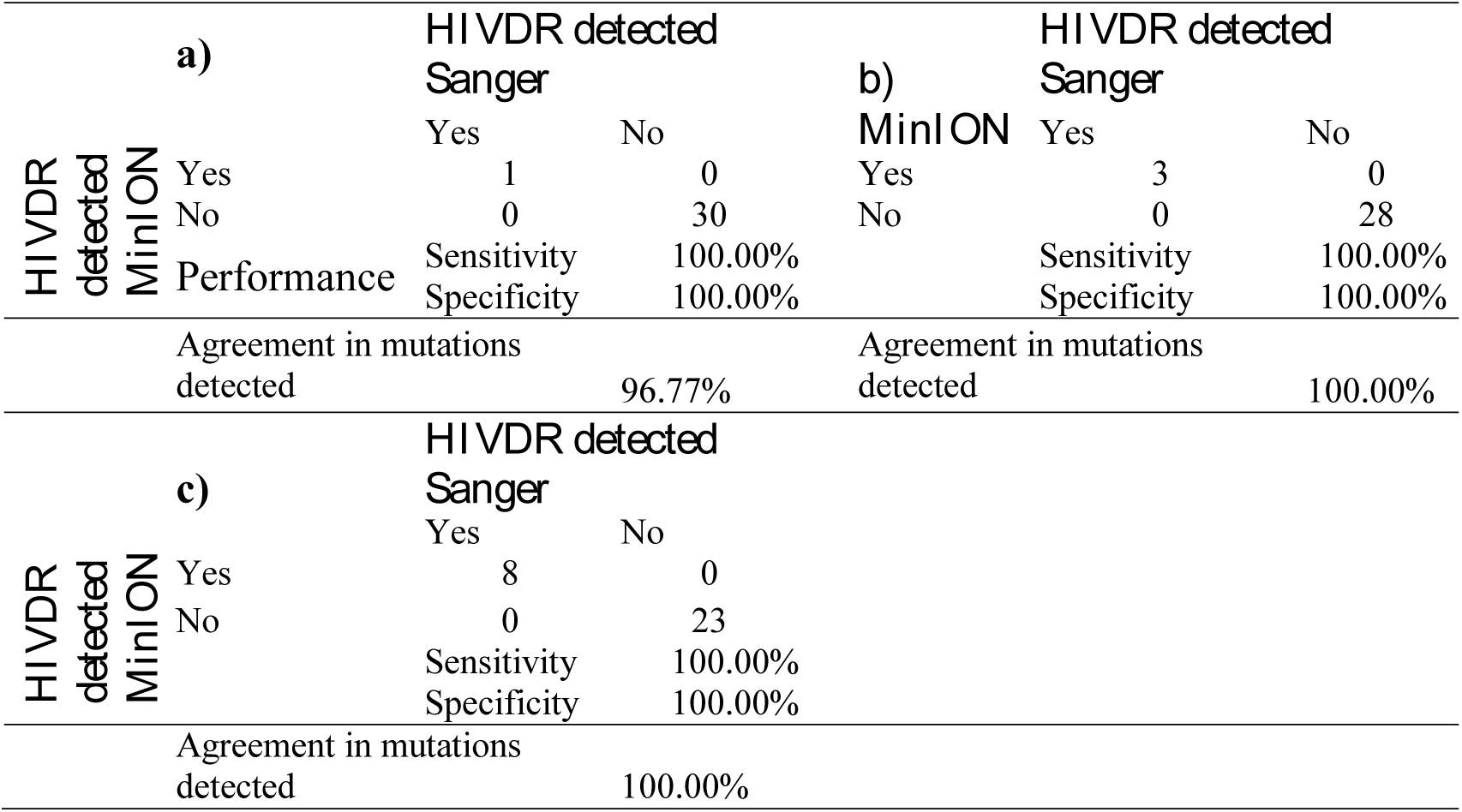
Evaluation of the Agreement of drug resistance interpretation in, Protease inhibitor (a), Nucleoside reverse transcriptase inhibitors (NRTIs) (b) and non-nucleoside reverse transcriptase inhibitors (NNRTIs) mutations (n=31)

Additionally, we found a good positive predictive value (100%) in the estimation of drug resistance mutations in the groups of protease inhibitors (PI), nucleoside reverse transcriptase inhibitors (NRTIs), and non-nucleoside reverse transcriptase inhibitors (NNRTIs), table #2. **Discussion**

The implementation of new methodologies for the analysis of HIV drug resistance with the capabilities of decentralization is a priority for health systems, specially in developing countries. Here we present a methodological approach for the sequencing and analysis of HIV sequences obtained with MinION Oxford Nanopore Sequencer.

The comparison of sequences generated with ONT and Sanger showed an excellent agreement with a percentage of identity greater than 95%. These results were obtained after the incorporation of a polishing process to the canu generated consensus, in which a draft generated from the reads was improved using Signal-level algorithms that harmonize the reads’s k-mers with the consensus generated with canu.

This approach to overcome the high error rate of ONT generated reads have been successfully used in other instances, as in the generation of the complete genome of *Escherichia coli* (9) and the de novo assemblies of four yeast strains (10). However, the weakness of this methodology is that the combination of canu-nanopolish required more total CPU time to complete the polishing, ∼2 hours/sample in a iCore 7, 16GB computer.

The analysis of the nucleotide distance provided good agreement between both methodologies compared, in addition, it was also good agreement in the interpretation of resistance when the HIV resistance database from Stanford University was used. These encouraging results pull forward to carry out studies with a wider number of samples. This will allow a more exhaustive validation of the clinical performance of this methodology in accordance with the WHO/ResNet guidelines (11).

Several studies have reported that setting up a NGS laboratory using MinION can be done without the use of expensive equipment and accessories (12, 13). Therefore, an HIV genotyping laboratory, based in ONT sequencing, can be implemented with standard equipment and well trained staff in molecular biology techniques. Our results strongly support the use of the MinION methodology as a viable option for decentralization of HIVDR testing in resource limited areas.

The method used to produce the HIV fragment for sequencing was previously implemented and used in several surveillance studies performed in our laboratory (8, 14). However, any method who efficiently amplify the region of interest (HIV pol: protease and RT genes) would produce similar results. The median length of the fragment obtained with the MinION agreed with the expected length of fragments used as input in the sequencing reaction, indicating that there was not production of PCR chimeras.

We observed a fast turnaround time for the MinION processing. Together with ARN extraction, PCR steps, MinION library preparation and sequencing, testing 12 samples can be completed in 2 days. However, using the Sanger method, 12 samples would take at least 12 more hours for processing when an ABI 3130xl 16 capillaries sequencer is used (supplementary Figure 1). Using barcode protocols, the cost per test with the MinION is almost the same than with the Sanger method (US$103) but it is very possible that the cost could be reduced for surveillance programs.

The ONT sequencing method has been experiencing many updates since its introduction in 2014. By the time this manuscript was written, there have been improvements in the chemistry an in the per-base reading accuracy. Additionally, some recent studies are showing implementation of more sophisticated bioinformatics tools for a better base calling of MinION generated reads (15).

Finally, we presented an approach for the analysis of HIV reads generated with MinION ONT, further studies are guaranteed before the application of this methodology in clinical settings.

## Acknowledgment

We would like to thank all the participants that agree to participate in the current study. This study was founded by grant FID14033 of the SENACYT-PANAMA. Alexander A. Martínez C. and Juan M. Pascale are members of the Sistema Nacional de Investigación (SNI) from SENACYT, Panama.

